# Extracts of food and medicinal plants sold in Moroccan markets induce apoptosis-like in *Toxoplasma gondii* tachyzoites *in vitro*

**DOI:** 10.1101/2024.07.19.603849

**Authors:** Ismail Elkoraichi, Nathalie Moiré, Samira Rais, Isabelle Dimier-Poisson, Fouad Daoudi, Françoise Debierre-Grockiego

## Abstract

Treatment of congenital toxoplasmosis is potentially toxic and above all too expensive to be administered systematically in middle-income countries such as Morocco. There is therefore a real interest in discovering alternative treatments that would be financially accessible to all. In this context, plants used in traditional medicine and purchased in markets are good candidates. Aqueous extract of *Ammi visnaga* seeds had no inhibitory effect against *T. gondii* tachyzoites intracellular growth *in vitro* and induced a cytotoxic effect on host cells. In contrast, ethanolic extract of *A. visnaga* seeds showed anti-*Toxoplasma* effect with low cytotoxicity, indicating the compounds extracted differed according to the solvent used. Aqueous extracts of *Punica granatum* peel and *Syzygium aromaticum* flower buds also showed anti-*Toxoplasma* effect with low cytotoxicity. All the extracts tested, with a greater effect from aqueous and ethanolic extracts of *A. visnaga* seeds, induced apoptosis-like of extracellular tachyzoites, as determined by exposure of phosphatidylserine on tachyzoite surface and DNA fragmentation. Finally, the aqueous extracts of *A. visnaga* seeds, *P. granatum* peel and *S. aromaticum* flower buds exhibited antioxidant properties. Phytochemical analysis indicated that coumarins and sterols from the aqueous extract of *A. visnaga*, saponins from the ethanolic extract of *A. visnaga*, gallic tannins from the extract of *P. granatum*, and phenols from the extract of *S. aromaticum* were certainly the main components responsible for the different effects observed. The results suggested that the seeds of *A. visnaga* and the peel of *P. granatum* are the two best candidates for possible preclinical studies on toxoplasmosis.

## Introduction

The unicellular eukaryotic parasite *Toxoplasma gondii* can cause severe damage to the eye (ocular toxoplasmosis), to the brain (cerebral toxoplasmosis) and to the foetus (congenital toxoplasmosis) with blindness, encephalitis and abortion as possible consequences. Toxoplasmosis is a ubiquitous infectious disease, but the prevalence is highly variable in different regions of the world with the highest prevalence on the African continent (61.4% calculated from 17 studies between 1995 and 2017) and the lowest prevalence on the Asian continent (16.4% calculated from 61 studies between 1996 and 2017) [34]. In addition to these mean values, there are large intracontinental variations: 20.8 – 87.7% for Africa and 0.5 – 82.6% for Asia [34]. In Morocco, overall human seroprevalence has been estimated to be between 40 and 60% [30]. Data on acute *T. gondii* infection during pregnancy indicated that the prevalence was 2.5% (1.7 – 3.4%) in the Eastern Mediterranean region, which includes Morocco according to the World Health Organisation classification, and 0.8% (0.3 – 1.5%) and 1% (0.5 – 1.6%) in the Western Pacific and in the South-East Asia regions, respectively [40]. Humans are infected by ingestion of oocysts disseminated in the environment within faeces of felids (definitive hosts) or by ingestion of cysts in undercooked meat (from intermediate hosts). Prevalence is therefore related to the number of infected felids, climate (temperature and humidity are parameters that play a role in the survival of oocysts in the environment), hygiene and income levels, and culinary habits. Regarding the influence of the climate in Morocco, some data indicated that the prevalence of severe congenital toxoplasmosis was higher in the Sahara region (8/10000 live births) than in the oceanic and mountainous regions (3.9 to 5.4/10000 live births) [16]. A more recent study using a low-cost rapid test conducted in different regions of Morocco on 632 women, 82 of whom were pregnant, showed that the prevalence of *T. gondii* infection was 2.5 to 3 times higher in women with a lower level of education living in mountainous and urban areas [17]. These populations were probably unaware of the risks and modes of infection of *T. gondii*. In the Maghreb countries, meat is usually highly cooked and most infections were suspected to be due to oocysts linked to infected definitive hosts. The seroprevalence of *T. gondii* was lower in domestic cats from the Asian continent [27% (24 – 30%)] compared to the African continent [51% (20 – 81%)], whereas the difference was not as dramatic for wild felids (67% in Asia and 74% in Africa) [36].

The current treatment of congenital toxoplasmosis is a combination of a 2,4-diaminopyrimidine (pyrimethamine, trimethoprim) and a sulfonamide (sulfadiazine, sulfamethoxazole) from the 18^th^ week of pregnancy. Spiramicin, suspected to be less effective against foetal transmission, but without the potent teratogenic effects of 2,4-diaminopyrimidine, is given during the first six months of pregnancy [32]. However, diagnosis of congenital toxoplasmosis followed by treatment is not routine in middle-income countries such as Morocco [16]. Thus, it would be useful to discover safe and effective anti-*Toxoplasma* activity in cheap products easily accessible to the population, such as plants used in traditional medicine and food. For several decades, plants have been tested for their effectiveness against *T. gondii*. They may be parts of trees, wild, medicinal or edible plants from all continents, with mainly aqueous and alcoholic extracts [41,42,13]. This wide interest may be explained by a good cultural acceptance of the populations already used to this kind of preparations and the belief that plant extracts have less side effects than chemical drugs. In the present work, the anti-*Toxoplasma* and the antioxidant effects of extracts of different plant parts freely sold on the markets of Moroccan cities were investigated: seeds of Khella/toothpick plant (*Ammi visnaga* L., Apiaceae), peel of pomegranate (*Punica granatum* L., Lythraceae) and flower buds of clove (*Syzygium aromaticum* L., Myrtaceae). *A. visnaga* is traditionally used to treat urological diseases, but an antimicrobial effect has also been demonstrated against fungi and bacteria [25]. *P. granatum* is known for its antioxidant and antimicrobial activities [18]. The essential oil or ethanolic extract of clove (*S. aromaticum*) has shown the highest antibacterial and antifungal activities among other spices such as basil, bay leaf, cinnamon, coriander, cumin, Jamaica pepper, lemongrass, oregano, rosemary and thyme [29]. *S. aromaticum* also has antioxidant, anti-inflammatory, anticancer, analgesic and anaesthetic properties [9]. Because of all these well-known applications in human and veterinary medicine, especially against microorganisms, these plants are good candidates for the study of their anti-*Toxoplasma* activity.

## Materials and methods

### Chemicals

Ascorbic acid, 2,2-diphenyl-1-picrylhydrazyl (DPPH), dimethyl sulfoxide (DMSO), hydrogen peroxide (H_2_O_2_), gallic acid, glucose, and quercetin were purchased from Sigma-Aldrich Chemicals. All solvents of analytical grade were purchased from Merck Limited.

### Plant collection and extraction

Khella/toothpick plant seeds (*Ammi visnaga* L., Apiaceae), pomegranate peel (*Punica granatum* L., Lythraceae) and clove flower buds (*Syzygium aromaticum* L., Myrtaceae) were purchased in the markets of Casablanca between June and September 2019. Their identification was done by Prof. Khayati Najat according to Bellakhdar [10].

Plant materials were washed with water, dried at room temperature and powdered. Aqueous extracts were prepared from the seeds of *A. visnaga* (Av-SA), the peel of *P. granatum* (Pg-PA) and the flower buds of *S. aromaticum* (Sa-FA) by adding 500 mL of water at 70 °C to 50 g of plant powder. The extractive solution was obtained by stirring until the water reached room temperature. Ethanolic extracts were prepared from *A. visnaga* seeds (Av-SE) by maceration of 50 g of plant powder with 500 mL of absolute ethanol for 72 h at room temperature in the dark. All preparations were then centrifuged at 2,000 g for 10 min, filtered through Whatman 3MM paper and dried using a vacuum rotary evaporator in a water bath at 55 °C. The extraction yield was calculated as (dry weight of extract/ dry weight of sample) x 100. Extracts were suspended in phosphate buffer saline (PBS) and stored at 4 °C.

### Growth inhibition of intracellular *T. gondii* tachyzoites

The efficacy of the plant extracts on *T. gondii* was tested in a colorimetric assay, as described previously [33]. The *T. gondii* strain RH-β-gal carrying the *E. coli lacZ* gene encoding β-galactosidase under the control of the *T. gondii sag1* promoter was used. Tachyzoites were maintained on human foreskin fibroblasts (HFF, ATCC number CRL-1634) cultivated in Dulbecco’s Modified Eagle Medium (DMEM), 2 mM L-glutamine and 10% fetal calf serum (FCS) at 37 °C in 5% CO_2_ atmosphere. For the assay, HFF were seeded in 96-well plates at 2 ×10^4^ cells/well in 100 µL of DMEM without phenol red (Pan Biotech GmbH), 2 mM L-glutamine and 1% FCS at 37 °C in 5% CO_2_ atmosphere. After 24 h, 100 RH-β-gal tachyzoites and plant extracts (50 - 15000 µg/mL in quadruplicate) were added to HFF at the same time, each in 50 µL of the same medium. The final ethanol concentration was well below 0.1%, which is known to be toxic to *T. gondii*. The medium alone (no parasite, no plant extract) was used as negative control. Pyrimethamine (Sigma) at 1 µM was used as a positive control to abrogate parasite development. After 96 h incubation at 37 °C in 5% CO_2_ atmosphere, β-galactosidase was released with a lysis solution (0.1% Triton X-100, Sigma) and its activity was measured by addition of 1 mM chlorophenol red-β-D-galactopyranoside (Roche Diagnostics, GmbH) in 100 mM HEPES pH 8 (Dutscher). The absorbance was read at 565 nm. The IC_50_ (50% inhibitory concentration) was calculated from the β-galactosidase activity in the presence of the plant extracts compared to untreated parasites.

### Cytotoxicity effect on host cells

HFF were cultured in 96-well plates at 2 ×10^4^ cells/well in 150 µL of DMEM with 2 mM L-glutamine, 1% fetal calf serum (FCS, Dominique Dutscher) and plant extracts at 50 - 1000 µg/mL in triplicate. Plates were incubated at 37 °C in a 5% CO_2_ atmosphere for 96 h. UptiBlue^TM^ (15 µL, Interchim) was then added for 4 h incubation in the dark at 37 °C in a 5% CO_2_ atmosphere. The absorbance was measured at 565 nm and 630 nm to calculate the reduction of the redox indicator. The CC_50_ (50% cytotoxic concentration) was calculated from the reduction in the presence of the plant extracts compared to untreated cells. The selectivity index was calculated by the ratio CC_50_/IC_50_.

### Killing activity on extracellular *T. gondii* tachyzoites

*A. T. gondii* tachyzoites were seeded in 24-well plates at 10^7^ in 2 ml DMEM with 2 mM L-glutamine and 1% FCS alone or with plant extracts at 1000 and 5000 µg/mL. After 96 h incubation at 37 °C in a 5% CO_2_ atmosphere, 10^5^ parasites were stained with an acridine orange solution (Sigma) at a final concentration of 0.005%. Dead (green cytoplasm and nucleus) and alive (orange cytoplasm and yellow nucleus) parasites were counted by microscopic observation under UV light (Olympus microscope).

### Phosphatidylserine exposure on extracellular *T. gondii* tachyzoites

*A. T. gondii* tachyzoites were seeded in 96-well plates at 2x10^5^ in 200 µL DMEM with 2 mM L-glutamine and 1% FCS alone or with plant extracts at concentrations from 0.5 to 500 µg/mL. After 24 h incubation at 37 °C in a 5% CO_2_ atmosphere, tachyzoites were stained with 2 µL of Annexin-V-FITC and 2 µL of propidium iodide (ab 14085, Abcam) in 100 µL annexin-V buffer (ab… Abcam) for 15 min at 4 °C in the dark. After centrifugation at 3,000 g, tachyzoites were suspended in annexin-V buffer and analysed with a Miltenyi cytometer. Data were exported with Flowlogic^®^ software (version 7.3).

### DNA fragmentation of extracellular *T. gondii* tachyzoites

*A. T. gondii* tachyzoites were seeded in 6-well plates at 2x10^7^ in 4 mL DMEM with 2 mM L-glutamine and 1% FCS alone or with plant extracts at concentrations from 50 to 5000 µg/mL. After 24 h incubation at 37 °C in a 5% CO_2_ atmosphere, tachyzoites were centrifuged at 3,000 g for 5 min at 4 °C and lysed by mixing the pellet with 1 mL of lysis buffer (10 mM Tris/HCl pH 8, 100 mM EDTA, 10 mM EGTA, 0.5% SDS), except 10^5^ tachyzoites that were used for determining exposure of phosphatidylserine as described above. Samples were incubated for 2 h at 37 °C with RNAse A at 30 µg/mL, followed by 1 h at 56 °C with proteinase K at 100 µg/mL. The lysates were then extracted with phenol-chloroform-isoamyl alcohol (25:24:1) by mixing for 10 min at 4 °C and centrifuged at 16,000 g for 15 min at 4 °C. The upper aqueous phase was divided in 2 tubes and treated each with 75 µL of 10 M sodium acetate and 900 µL of absolute ethanol. DNA was precipitated at -20 °C for 20 min and centrifuged at 16,000 g for 15 min at 4 °C. Supernatants were discarded and the two pellets of each sample were pooled in 1 mL 70% ethanol. DNA was spun at 16,000 g for 1 min at 4 °C, air dried and dissolved in 30 µL of 5 mM Tris/HCl PH 8.5 for 5 min at 65 °C.

### Antioxidanht activity

The antioxidant activity of the plant extracts was measured by DPPH (2,2-diphenyl-1-picrylhydrazyl, Sigma-Aldrich) radical scavenging activity tests [21]. Twenty µL of plant extracts at different concentrations were added to 180 µL of 60 µM DPPH. Absorbance was read at 517 nm after 30 min incubation at room temperature in the dark. The percentage of DPPH free radical scavenging was calculated as ([absorbance of blank/absorbance of plant extract]/absorbance of blank) x 100, with methanol as the blank. Ascorbic acid (Sigma-Aldrich) was used as positive control for its antioxidant property.

### Phytochemical screening

Semi-quantitative methods were used to identify the different classes of active chemical constituents of the plant extracts [27]. All analytical grade solvents were purchased from Merck Limited. The presence of saponins was detected by the formation of a foam persisting at least 15 min after vigorously mixing 1 volume of the plant extract with 2 volumes of water. To detect the presence of sterols and triterpenes, the dried plant extracts were dissolved in 1 volume of acetic anhydride, 1 volume of chloroform and 4 volumes of sulphuric acid and the appearance of a purple and then blue and green ring indicated a positive reaction. A yellow precipitation after addition of Mayer’s reagent (potassium mercuric iodide solution) to plant extracts diluted in 1% hydrochloric acid indicated the presence of alkaloids. The yellowish-white colour after addition of 10% NaOH to plant extracts indicated the presence of coumarins. Dark blue and green precipitation after addition of 3 drops of 1% ferric chloride solution indicated the presence of gallic tannins and catechol-type tannins, respectively. The red-orange colour after the addition of metallic magnesium and concentrated hydrochloric acid to the heated plant extracts indicated the presence of flavonoids (Shinoda test). The Folin-Ciocalteu method was used to detect the presence of phenols in plant extracts dissolved at 1 mg/mL in methanol, given a blue-green colour [4]. Addition of aluminium chloride to the plant extract led to the appearance of a yellow colour in the presence of flavonoids and the addition of aluminium chloride plus sodium acetate changed the extract to a dark yellow colour in the presence of flavonols.

Phenols and flavonoids contents were also calculated as mg/g of gallic acid and quercetin equivalents, respectively, from calibration curves.

### Statistical analysis and figures

Statistical analyses were done with Prism 7 (GraphPad software, Inc.) using the one-way ANOVA test followed by Sidak’s multiple comparisons test or the Kruskal-Wallis test followed by Dunn’s multiple comparison test. Figures were produced with the same software.

## Results

### Effect of plant extracts on intracellular growth of *T. gondii* and on host cell cytotoxicity

The preparation of aqueous plant extracts has the advantage of not requiring expensive and polluting chemicals. In this study, *T. gondii* tachyzoites and aqueous extracts of *A. visnaga*, *P. granatum* and *S. aromaticum* were simultaneously added to HFF and incubated for 96 h so that active molecules could act on several lytic cycles of the parasite, from the invasion stage to the egress stage. As is usually the case with products containing many different compounds [15], the concentration inducing a 50% reduction in parasite numbers (IC_50_) was variable from experiment to experiment, resulting in high standard deviations. Nevertheless, the aqueous extract of *A. visnaga* seeds was the least effective against *T. gondii* with an IC_50_ of 679.6 ± 243.6 µg/mL (Table 1). Aqueous extracts of *P. granatum* peel and *S. aromaticum* flower buds showed relatively good efficacy with an IC_50_ around 400-500 µg/mL. To understand to what extent the extraction method could play a role in the antiparasitic effect, the seeds of *A. visnaga* were extracted with ethanol. This extract was the most effective against *T. gondii* with an IC_50_ of 192.3 ± 70.2 µg/mL.

**Table 1.** Effect of plant extracts on *T. gondii* growth and cytotoxicity on HFF.

|  | IC<sub>50</sub> (µg/mL) | CC<sub>50</sub> (µg/mL) | SI |
| --- | --- | --- | --- |
| Av-SA | 679.6 ± 243.6 | 313.3 ± 357.6 | 0.5 |
| Av-SE | 192.3 ± 70.2 | 515.9 ± 312.0 | 2.7 |
| Pg-PA | 413.5 ± 271.4 | 1007.4 ± 286.3 | 2.4 |
| Sa-FA | 473.5 ± 196.0 | 985.6 ± 388.6 | 2.1 |

Values are means ± standard deviations of three (Av-SA, Av-SE, Pg-PA) or four (Sa-FA) experiments with the same batch of plant extracts.
Abbreviations: IC<sub>50</sub>, 50% inhibitory concentration on *T. gondii* tachyzoite growth; CC<sub>50</sub>, 50% cytotoxic concentration on HFF; SI, selectivity index (CC<sub>50</sub>/IC<sub>50</sub>), Av-SA, *Ammi visnaga* – Seeds – Aqueous extract; Av-SE, *Ammi visnaga* – Seeds – Ethanolic extract; Pg-PA, *Punica granatum* – Peel – Aqueous extract; Sa-FA, *Syzygium aromaticum* – Flower buds – Aqueous extract.

Toxicity of the extracts was estimated on HFF, the *T. gondii* host cells used here, by calculating the concentration leading to 50% cytotoxicity compared to the untreated condition (Table 1). The aqueous extract of *P. granatum* peel was the least toxic to HFF (CC_50_ = 1007.4 ± 286.3 µg/mL), while the aqueous extract of *A. visnaga* seeds was the most cytotoxic (CC_50_ = 313.3 ± 357.6 µg/mL).

A high selectivity index calculated with both values (SI = CC_50_/IC_50_) is an indicator of the good benefit-risk balance of a treatment. The selectivity index was greater than 2 for *A. visnaga* seed ethanolic extract, *P. granatum* peel aqueous extract and *S. aromaticum* flower bud aqueous extract, but less than 1 for *A. visnaga* seed aqueous extract due to its low anti-*Toxoplasma* activity (Table 1).

### Killing effect of plant extracts on extracellular *T. gondii* tachyzoites

To understand whether the cytotoxicity towards HFF reflects a direct killing effect of plant extracts on *T. gondii*, free tachyzoites were incubated for 96 h with the aqueous extract of *A. visnaga* seeds or of *P. granatum* peel at 1000 and 5000 µg/mL, i.e. from 1.4 to 12 times the IC_50_ for 10^5^ times more parasites. Dead parasites (Fig. 1A, top picture) were distinguished from live parasites (Fig. 1A, bottom picture) by a different colour (green *vs*. yellow-orange) after staining with acridine orange. Compared to the control condition (medium alone), the *A. visnaga* extract did not increase the percentage of tachyzoite mortality at 1000 µg/mL, whereas it led to 100% death at 5000 µg/mL (Fig. 1B). In contrast, *P. granatum* extract had no killing effect on *T. gondii* tachyzoites at 1000 µg/mL and only a moderated increase in the percentage of tachyzoite mortality at 5000 µg/mL. Thus, the most cytotoxic extract on HFF was lethal to *T. gondii* and the least cytotoxic extract on HFF was not lethal to the parasite.

**Figure 1.**
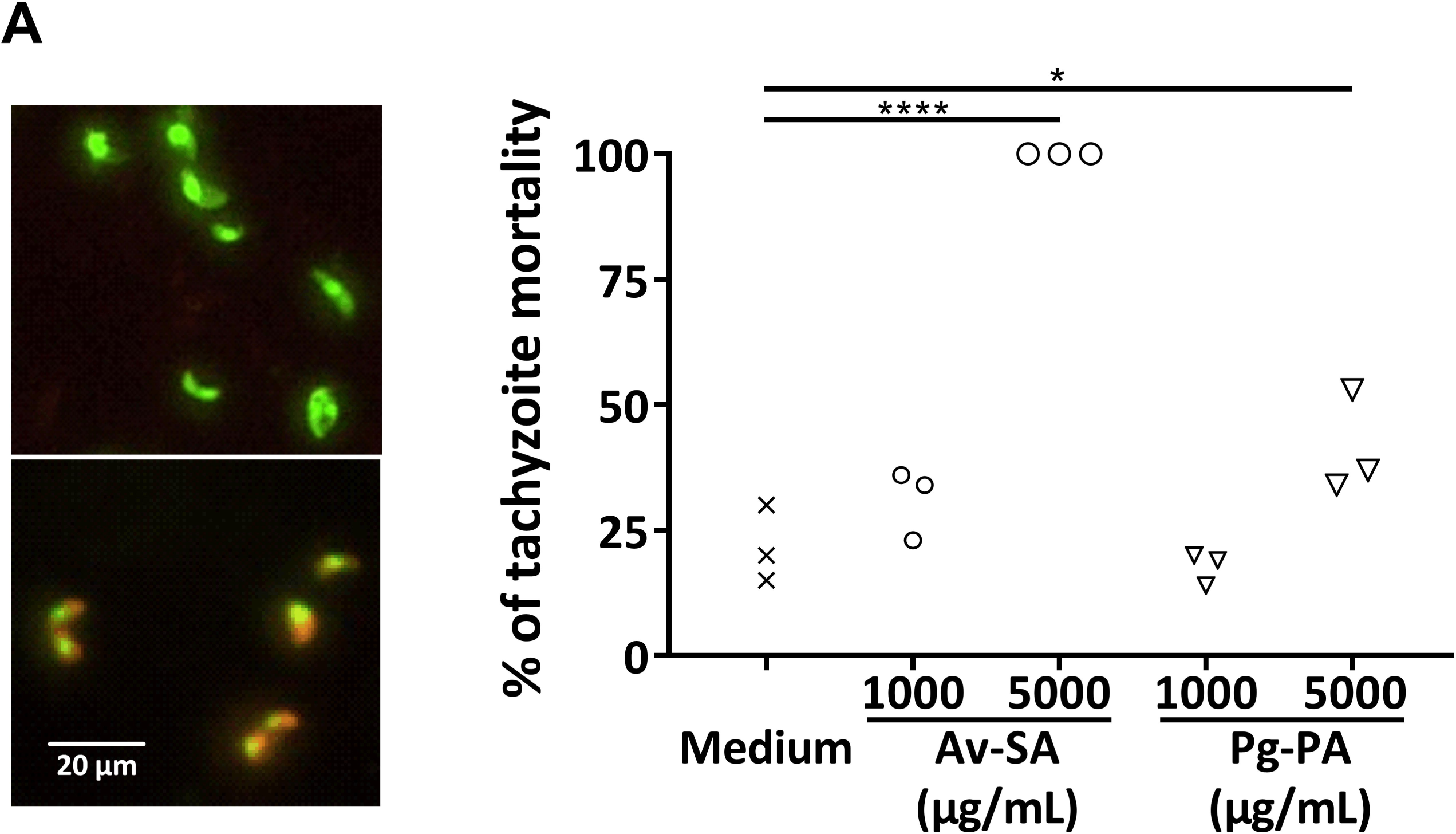
Killing activity of two plant extracts. Mortality of *T. gondii* was determined after incubation for 96 h of free tachyzoites with medium alone or with the plant extracts at 1000 and 5000 mg/mL followed by their staining for 30 min with acridine orange, allowing to distinguish dead (**A**, upper picture) and living (**A**, lower picture) tachyzoites, scale bar 20 µm. (**B**): The values represent the percentage of dead tachyzoites from 100 total tachyzoites counted on 3 microscopic fields of one experiment. **** p<0.0001 and * p<0.05 with the Sidak’s multiple comparisons test. Av-SA: *Ammi visnaga* – Seeds – Aqueous extract; Pg-PA: *Punica granatum* – Peel – Aqueous extract.

### Apoptosis-like induced by plant extracts on extracellular *T. gondii* tachyzoites

After this preliminary experiment with the maximal concentrations tested in this study, apoptosis traits were explored to understand whether these plant extracts could induce this type of death for *T. gondii* tachyzoites in a dose-dependent manner. Exposure of phosphatidylserine at the membrane surface of tachyzoites treated for 24 h with 0.5 to 500 µg/mL of plant extracts was quantified by flow cytometry after labelling with annexin V-FITC and propidium iodide. The tachyzoite population did not differ in size or density after treatment at all concentrations tested of the 4 extracts. This is illustrated in Figure 2 with the ethanolic extract of *A. visnaga* (Fig. 2 A, C, E, G, I). In addition, the treatment increased the percentage of annexin V+ tachyzoites, while the percentage of propidium iodide+ tachyzoites remained below 5% (Fig. 2 B, D, F, H, J), indicating death by apoptosis rather than necrosis. All 4 plant extracts increased the percentage of apoptotic tachyzoites at the lowest concentration tested (0.5 µg/mL), with a significant difference for the aqueous extract of *A. visnaga* seeds (Av-SA) compared to the untreated condition (Fig. 3). The percentage of annexin V+ tachyzoites decreased at 5 µg/mL and 50 µg/mL and increased again at 500 µg/mL with all aqueous extracts (p<0.01 for Av-SA at 500 µg/mL), whereas it increased at 5 µg/mL and 50 µg/mL (p<0.01) and decreased at 500 µg/mL with the ethanolic extract of *A. visnaga* (Av-SE). This again shows that the solvent used to extract the active compounds has a significant impact on the activities of the plant extracts. Microscopic observation revealed an aggregation of tachyzoites in the presence of plant extracts, which could render their membrane inaccessible to annexin V (not shown).

**Figure 2.**
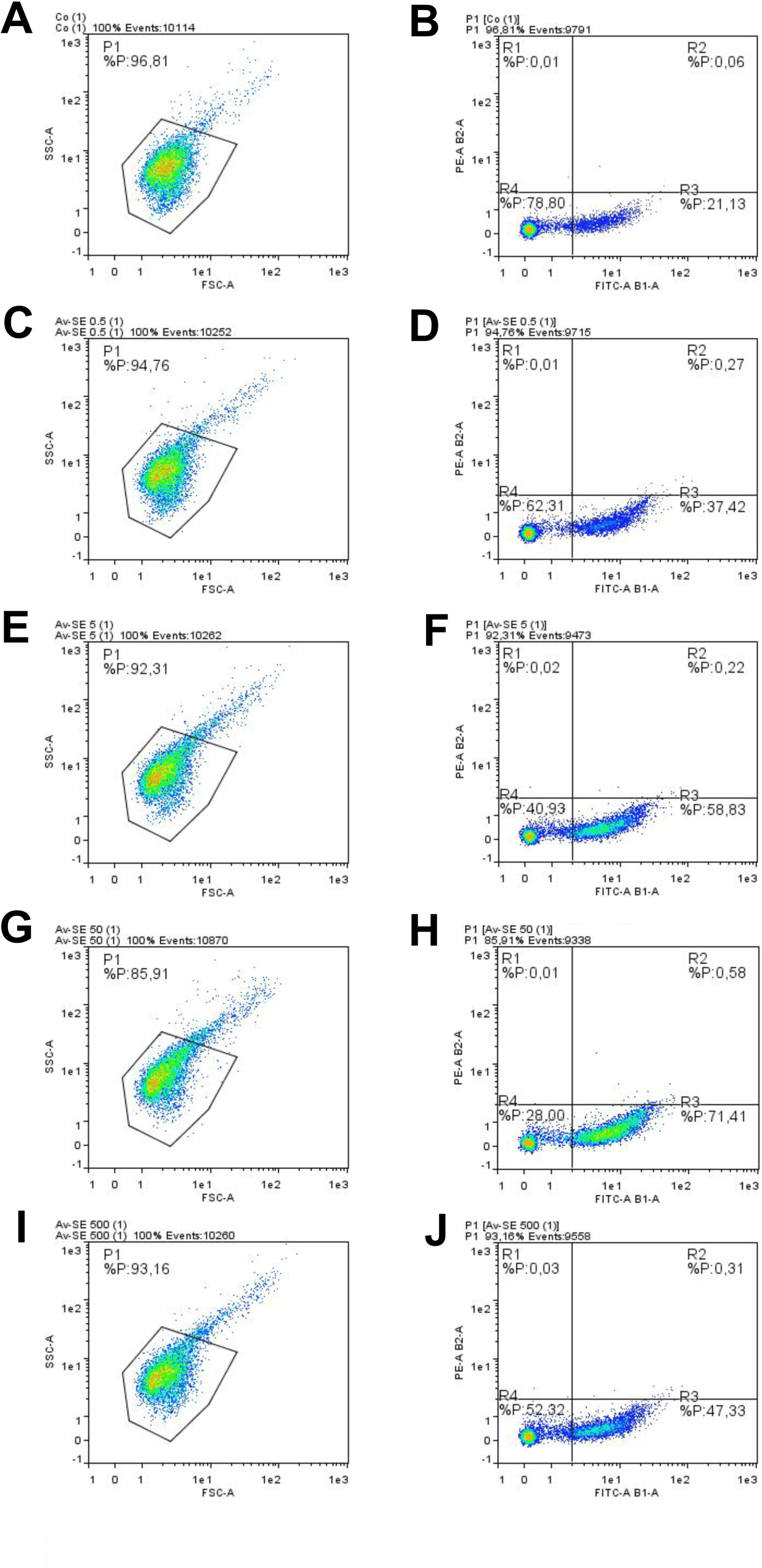
Phosphatidylserine exposure induced by *A. visnaga* seed ethanolic extract. *T. gondii* tachyzoites were incubated for 24 h with medium (**A**, **B**) or with *A. visnaga* seed ethanolic extract (Av-SE) at 0.5 µg/mL (**C**, **D**), 5 µg/mL (**E**, **F**), 50 µg/mL (**G**, **H**), or 500 µg/mL (**I**, **J**). Tachyzoites were labelled with annexin V-FITC and propidium iodide and analysed by flow cytometry. (**A**, **C**, **E**, **G**, **I**) Distribution of total and P1-gated populations according to forward scatter (FCS) and side scatter (SSC); (**B**, **D**, **F**, **H**, **J**) Annexin V-FITC and propidium iodide fluorescence intensity of P1 population. R2: annexin V+/propidium iodide+ (necrotic) tachyzoites; R3: annexin V+/propidium iodide-(apoptotic) tachyzoites; R4 : annexin V-/propidium iodide-(living) tachyzoites.

**Figure 3.**
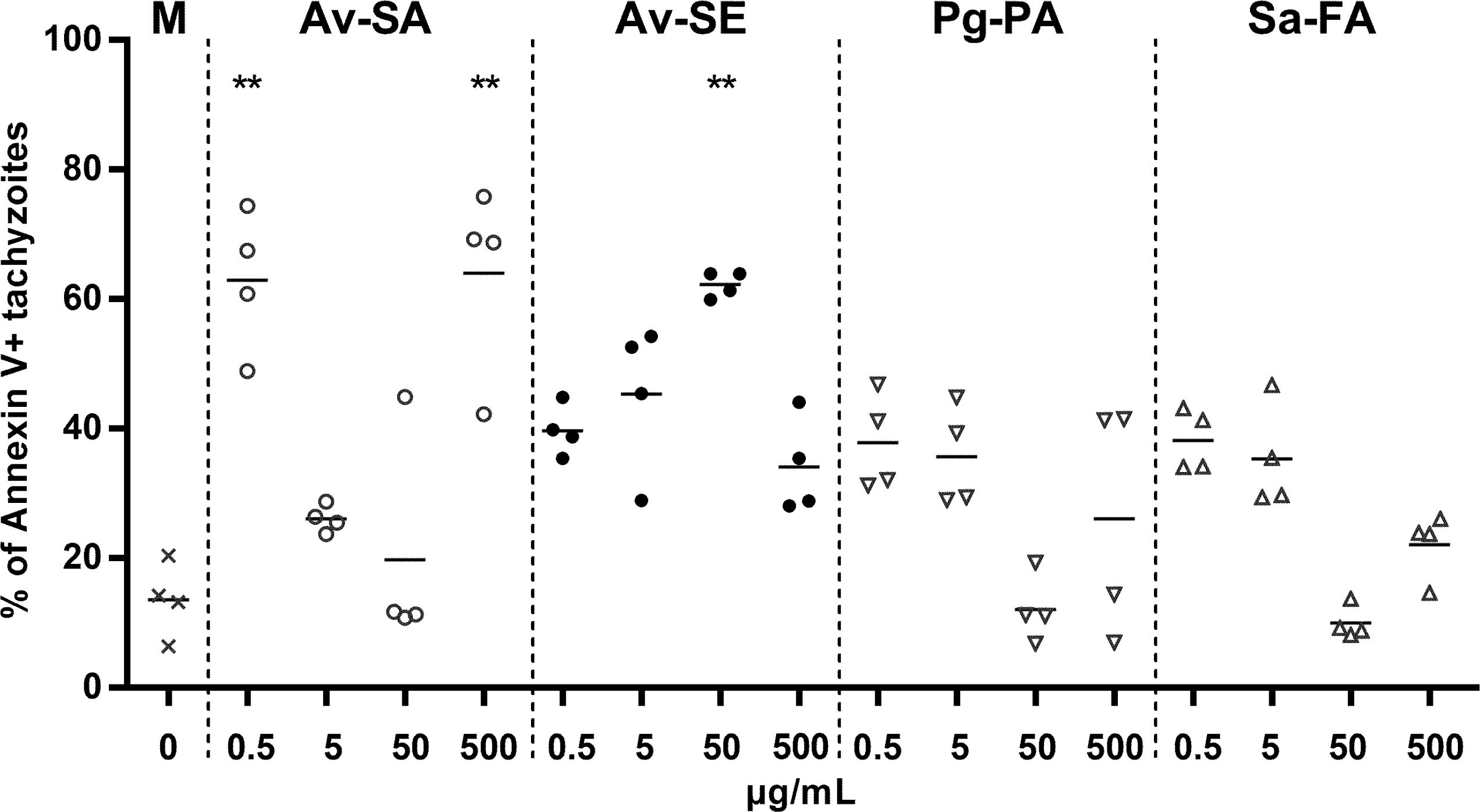
Phosphatidylserine exposure induced by plant extracts. *T. gondii* tachyzoites were incubated for 24 h with medium alone or with plant extracts at 0.5, 5, 50, or 500 µg/mL. Tachyzoites were labelled with annexin V-FITC and propidium iodide to analyse phosphatidylserine exposure by flow cytometry. Data represent individual values and median of annexin V+/propidium iodide-tachyzoites of 4 experiments. ** p<0.01 with the Dunn’s multiple comparisons test. M: medium alone; Av-SA: *Ammi visnaga* – Seeds – Aqueous extract; Av-SE: *Ammi visnaga* – Seeds – Ethanolic extract; Pg-PA: *Punica granatum* – Peel – Aqueous extract; Sa-FA: *Syzygium aromaticum* – Flower buds – Aqueous extract.

Visualisation of DNA fragmentation was then used to confirm that the plant extracts induced apoptosis-like in *T. gondii* extracellular tachyzoites. After incubation for 24 h, the DNA was purified and separated by electrophoresis. At the same time, phosphatidylserine exposure was quantified by cytometry. For conditions with 50 µg/mL aqueous extract of *A. visnaga* seeds and 50 and 500 µg/mL aqueous extract of *P. granatum* peel, the percentage of annexin V+ tachyzoites was between 4% and 11% and no fragmentation was observed (Fig. 4). For conditions with 500 µg/mL aqueous extract of *A. visnaga* seeds and 5000 µg/mL aqueous extract of *P. granatum* peel, the percentage of annexin V+ tachyzoites was 17% and 26%, respectively, and DNA fragmentation was observed. In contrast, the aqueous extract of *S. aromaticum* flower buds triggered DNA fragmentation at all concentrations tested, even when only 9% of the tachyzoites were annexin V+ (Fig. 4). Thus, a direct relationship can be established between phosphatidylserine exposure and DNA fragmentation for aqueous extracts of *A. visnaga* seeds and *P. granatum* peel, but not for that of *S. aromaticum* flower buds.

**Figure 4.**
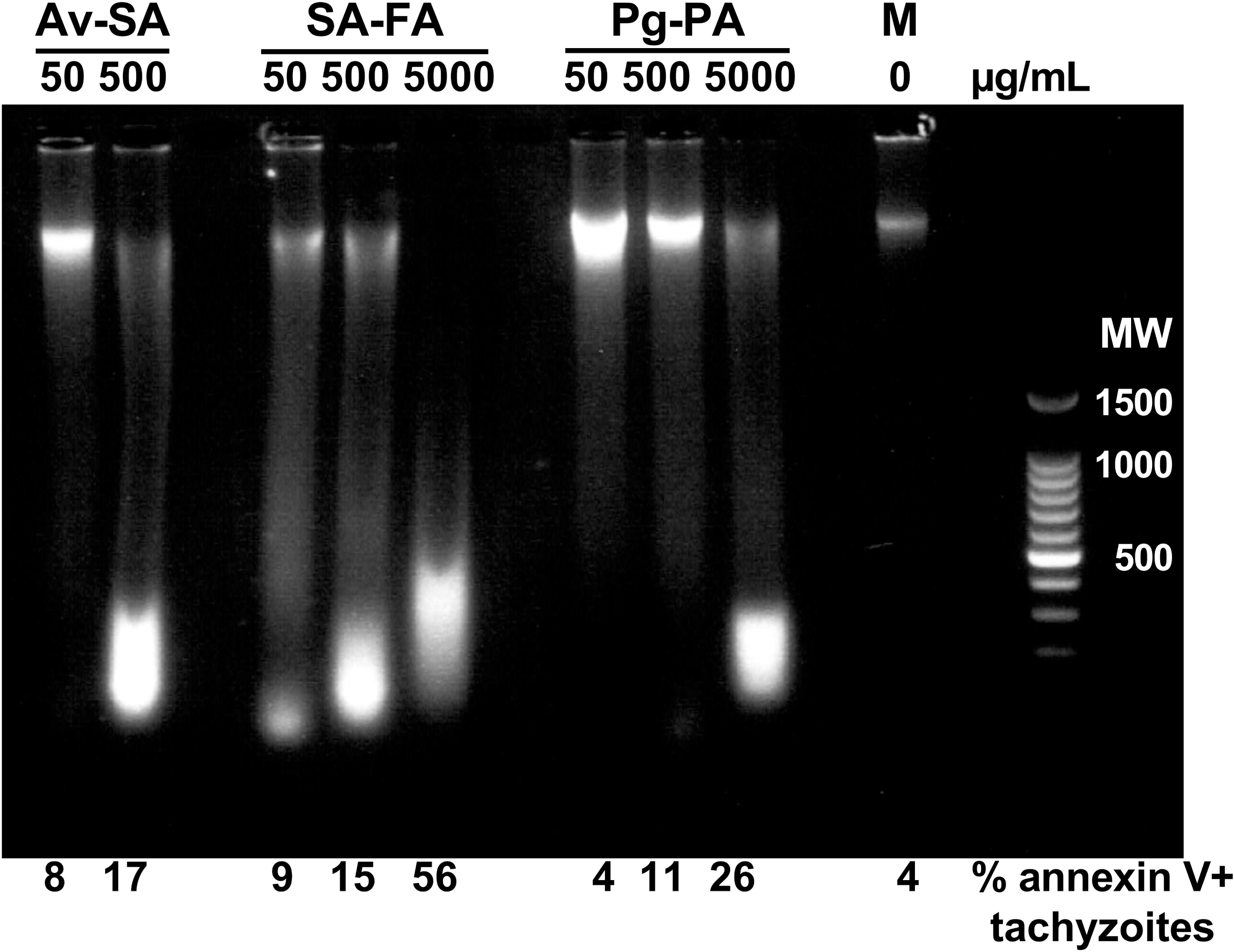
DNA fragmentation induced by plant extracts. *T. gondii* tachyzoites were incubated for 24 h with medium alone or with plant extracts at 50, 500 or 5000 µg/mL. Tachyzoite DNA was extracted and submitted to electrophoresis to visualise its fragmentation. Some tachyzoites were used to analyse phosphatidylserine exposure by flow cytometry (percentages of annexin V+ tachyzoites are indicated below the picture). Av-SA: *Ammi visnaga* – Seeds – Aqueous extract; Sa-FA: *Syzygium aromaticum* – Flower buds – Aqueous extract; Pg-PA: *Punica granatum* – Peel – Aqueous extract; M: medium alone; MW: molecular weight (100bp DNA ladder, Promega).

### Antioxidant properties of the plant extracts

The antioxidant activity of the plant extracts was assessed by the DPPH assay (Fig. 5). Ascorbic acid, used as a positive control, induced about 80% inhibition of DPPH oxidation in the range of 50 to 400 µg/mL. The aqueous extract of *A. visnaga* seeds, which had a selective index of 0.5, showed a linear dose-response to achieve the same efficacy as ascorbic acid at 400 µg/mL (Fig. 5). In contrast, the ethanolic extract of *A. visnaga*, which had the highest selectivity index (2.7), had negligible antioxidant activity with 20% inhibition of oxidation at 400 µg/mL, which is lower than the percentage obtained with 10 µg/mL ascorbic acid. Thus, the aqueous extract of *P. granatum* peel, which also had a good selectivity index (2.4), was tested in the DPPH trial. This extract had similar antioxidant activity to the aqueous extract of *A. visnaga* (Fig. 5). Finally, the aqueous extract of *S. aromaticum* flower buds showed lower antioxidant activity than the aqueous extract of *A. visnaga* between 50 and 200 µg/mL, but showed similar activity at 400 µg/mL.

**Figure 5.**
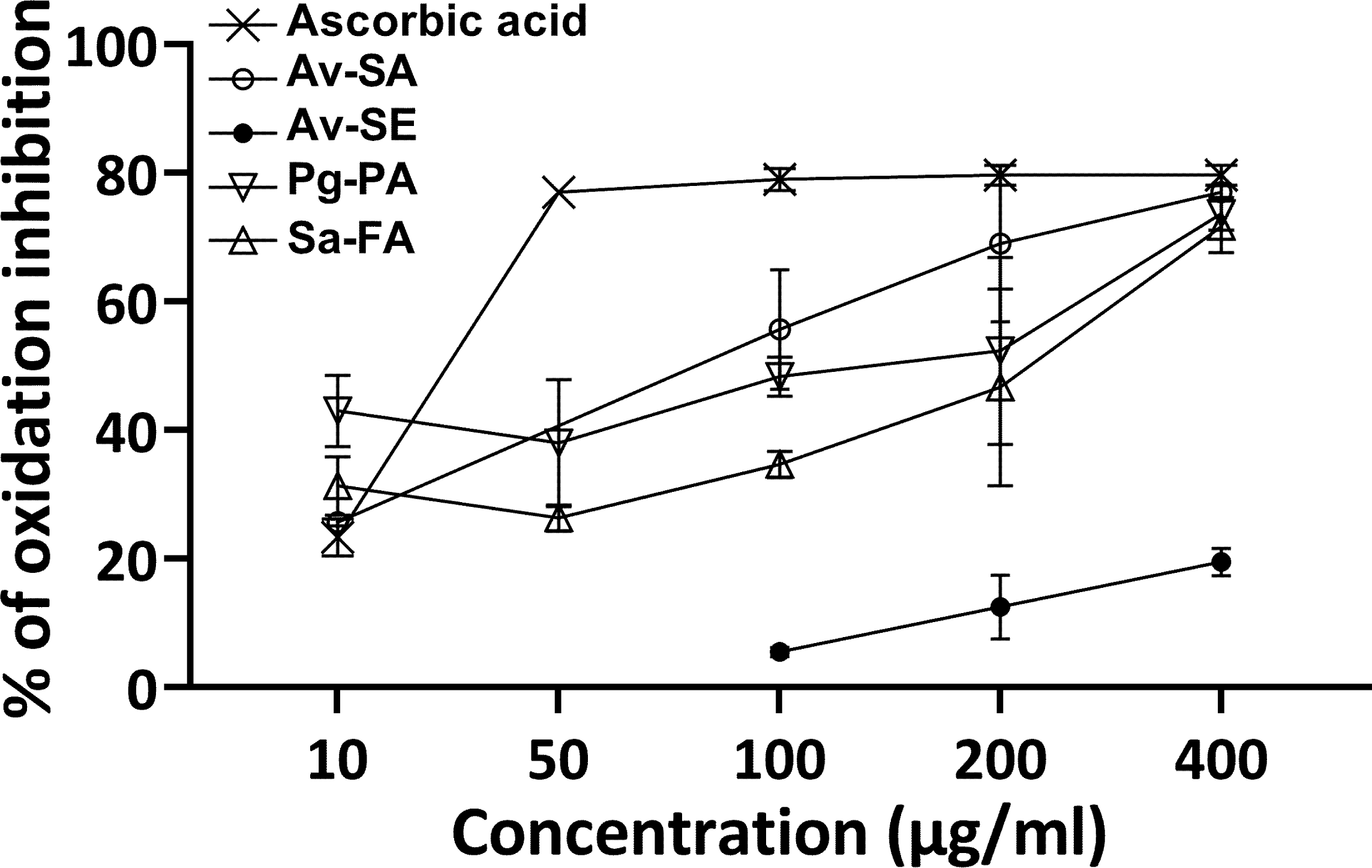
Antioxidant activity of the plant extracts. Scavenging of radicals from DPPH by the different plant extracts at concentrations between 10 and 400 µg/mL was measured at 517 nm after 30 min incubation at room temperature in the dark. Values are expressed as means ± standard deviations of the percentage of oxidation inhibition of three independent experiments with the same batch of plant extracts. Av-SA: *Ammi visnaga* – Seed – Aqueous extract; Av-SE: *Ammi visnaga* – Seed – Ethanolic extract; Pg- PA: *Punica granatum* – Peel – Aqueous extract; Sa-FA: *Syzygium aromaticum* – Flower buds – Aqueous extract.

### Phytochemical screening of the plant extracts

To understand which compounds extracted from the plants could be responsible for the different biological activities, the main constituents were identified by phytochemical analysis (Table 2). The aqueous extract of *A. visnaga* seeds had a low selectivity index (0.5) and antioxidant effects. This extract was rich in coumarins and sterols with intermediate amounts of phenols, saponins and cathechol tannins. The ethanolic extract of *A. visnaga* seeds, which showed the best selectivity index (2.7), but no antioxidant activity, contained mainly saponins, whereas phenols, coumarins, cathechol tannins and sterols were in intermediate amounts. Thus, the main difference with the aqueous extract of *A. visnaga* is the class of saponin compounds, which could have anti-*Toxoplasma* activity. The aqueous extract of *P. granatum* peel, which had the second highest selectivity index (2.4) due to lack of cytotoxicity and good anti-*Toxoplasma* activity, contained high amounts of phenols, gallic tannins and sterols. The aqueous extract of *S. aromaticum* flower buds (SI = 2.1) mainly contained phenols, flavonols and coumarins. Overall, these results indicate that the phytochemicals involved in anti-*Toxoplasma* and antioxidant activity were probably different and that the extraction method used plays an important role in the effect of the extracts.

**Table 2.** Phytochemical screening of the plant extracts.

|  | Av-SA | Av-SE | Pg-PA | Sa-FA |
| --- | --- | --- | --- | --- |
| Alkaloids | - | - | - | - |
| Catechol tannins | ++ | ++ | - | - |
| Coumarins | +++ | ++ | ++ | +++ |
| Flavonoids | + | + | + | ++ |
| Flavonoids (mg QE/g) | 0.054 ± 0.02 | 0.015 ± 0.02 | 0.008 ± 0.03 |  |
| Flavonols | - | - | - | +++ |
| Gallic tannins | - | - | +++ | + |
| Phenols | ++ | + | +++ | +++ |
| Phenols (mg GAE/g) | 0.725 ± 0.15 | 0.247 ± 0.012 | 1.368 ± 0.19 |  |
| Saponins | ++ | +++ | + | - |
| Sterols | +++ | ++ | +++ | + |
| Triterpenes | - | - | - | - |
| Extraction yield (%) | 47 | 12 | 56 | 30 |

Av-SA: *Ammi visnaga* – Seed – Aqueous extract; Av-SE: *Ammi visnaga* – Seed – Ethanolic extract; Pg-PA: *Punica granatum* – Peel – Aqueous extract; Sa-FA: *Syzygium aromaticum* – Flower buds – Aqueous extract; -, +, ++, +++ increasing amounts of compounds; mg QE/g: mg of quercetin equivalents/g of plant extract (mean ± standard deviation of 3 quantifications of the same batch); mg GAE/g: mg of gallic acid equivalents/g of plant extract (mean ± standard deviation of 3 quantifications of the same batch).

## Discussion

Selectivity index is an indicator determining whether a product could be a good antimicrobial candidate. For *T. gondii*, the selectivity index takes into account two distinct parameters, antiparasitic effect and cytotoxicity against host cells. Pyrimethamine has variable selectivity index towards *T. gondii* depending on the cells and study: either 0.9 and 0.89 using HeLa cells [43,19] or 23.8 and > 150 using HFF [1,33]. However, pyrimethamine is not recommended during the first trimester of pregnancy because of its potent teratogenic effect. This highlights the need for alternative safe treatment and additional toxicity testing before its administration to pregnant women.

In the present study, plant extracts were analysed for their anti-*Toxoplasma* activity, and in particular for the apoptosis-like effect induced in extracellular tachyzoites. In the literature, very few studies have investigated the mechanisms involved in apoptosis-like of *T. gondii* tachyzoites. Molecules known to be involved in apoptosis pathways were identified using a bioinformatics approach: a Tudor Staphylococcal Nuclease (substrate of apoptosis proteases), homologs of DNases (Endo G and ZEN1, but not CAD, ICAD and NUC1) and of Apoptosis Induction Factor (involved in nuclear apoptosis) [23]. In a 2013 publication, it was shown that treatment with staurosporine or miltefosine increased the percentage of annexin V+ tachyzoites, with no change in the percentage of 7-AAD+ tachyzoites, a marker of necrosis, and increased the percentage of TUNEL+ tachyzoites, a marker of DNA strand breaks [38]. Interestingly, antiparasitic drugs (anisomycin, atovaquone, clindamycin, pyrimethamine) were also able to increase the percentage of TUNEL+ tachyzoites, with the strongest effect being obtained with atovaquone. After treating extracellular tachyzoites for 1 h at 30 µg/mL, an aqueous extract of *Moringa oleifera* seeds increased the percentage of both Annexin V+/propidium iodide- and Annexin V+/propidium iodide+ parasite subpopulations. Higher concentrations were not evaluated against *T. gondii* because the viability of HeLa cells was reduced by 50% at 100 µg/mL [39].

The aqueous extract of *A. visnaga* seeds was antioxidant, but cytotoxic and no efficient against intracellular *T. gondii* tachyzoites (selectivity index = 0.5), whereas ethanolic extract of *A. visnaga* was not antioxidant, but efficient against intracellular *T. gondii* tachyzoites and not cytotoxic (selectivity index = 2.7). This ethanolic extract showed a direct dose-response apoptotic effect on extracellular tachyzoites. Information on the toxicity of *A. visnaga* extracts comes mainly from studies on their anticancer properties. For example, an hydro-ethanolic extract of *A. visnaga* fruits/seeds had a strong antitumor effect on Hep-G2 (liver carcinoma) cell line with a CC_50_ at 16.3 µg/mL [7]. In contrast, a methanolic extract showed no activity against Hela (cervical carcinoma cell line), but mild antitumor activity against Hep-G2 (liver carcinoma), HCT 116 (colon carcinoma) and MCF7 (breast carcinoma) cell lines with CC_50_ values ranging from 88 ± 1.99 μg/ml to 97 ± 2.05 μg/ml [11]. Thus, CC_50_ values were much lower on the different malignant cell lines tested in the literature than on HFF in our culture conditions. *In vivo*, *A. visnaga* was considered highly toxic on larvae of the invertebrate species *Artemia salina* (brine shrimp) with a concentration required for 50% lethality of 8.12 µg/mL for the ethanolic extract and 32.63 µg/mL for the aqueous fruits/seeds extract [20]. However, oral gavage of rats with 600 mg/kg/day of a hydro-ethanolic extract of *A. visnaga* seeds for 28 days induced no mortality and no changes in the levels of haematological, hormonal, renal and hepatic markers [26]. Thus, toxicity depends on the type of *A. visnaga* extract. The coumarin compounds khellin and visnagin purified from a methanolic extract of *A. visnaga* fruits/seeds had cytotoxic activity against cancer cells [11] and visnagin directly isolated from *A. visnaga* fruits/seeds was able to inhibit the growth of malignant cells through the induction of intracellular reactive oxygen species and apoptosis [8]. Thus, these two molecules could be responsible for the cytotoxic effect on HFF and the direct killing effect on *T. gondii* tachyzoites of the aqueous extract of *A. visnaga* seeds observed in our study.

In other studies using the DPPH assay, 50% inhibition of oxidation was obtained with 36 µg/mL aqueous extract of *A. visnaga* fruits/seeds, with 41 µg/mL ethanolic extract of *A. visnaga* fruits/seeds or with 50 µg/mL butanolic extract of *A. visnaga* aerial parts [20,12]. The authors suggested that the antioxidant activity of *A. visnaga* aerial parts was certainly due to the high content of flavonols, in particular quercetin [12]. Flavonols could not be detected in our aqueous extract of *A. visnaga* seeds and other components were certainly responsible for its antioxidant activity.

We have also found that aqueous extract of pomegranate peel was antioxidant, efficient against *T. gondii* and not cytotoxic (selectivity index = 2.4). An ethanolic extract of *P. granatum* fruit peel had a CC_50_ of 16.1 µg/mL on HFF and an IC_50_ of 82.8 on a RH-GFP strain of *T. gondii*, given a selectivity index of 0.2 [28]. The discrepancy with our result could be due to the differences in bioactive compounds extracted in water and ethanol. An aqueous extract of *P. granatum* peel was successfully used to treat cryptosporidiosis *in vivo*, leading to the absence of *Cryptosporidum parvum* oocysts in the faeces of mice orally treated with 3 g/kg for 3 days from 7 days post-infection [6]. Interestingly, treatment of chicks with 100-400 mg/kg of a methanolic extract of *P. granatum* peel completely stopped excretion of *Eimeria tenella* oocysts in faeces, associated with improved growth [2]. Similarly, a hydro-methanolic extract of *P. granatum* peel was as effective as ivermectin and albendazole in eliminating parasitic gastroenteritis eggs in faeces of different ruminant species after 2 oral treatments at 200 mg/kg [24]. These results indicate that the activity of the plant is not only dependent on the type of extract, but also on the target species. Although large quantities of *P. granatum* fruits can be ingested without major complications, the toxicity of *P. granatum* peel needs to be better documented. *In vitro*, acetonic, ethanolic and aqueous extracts of *P. granatum* peel showed less than 20% inhibition of viability on Chang liver and Vero kidney cell lines at 100 µg/mL [5]. *In vivo*, a single dose of 2000 mg/kg of a methanolic extract of *P. granatum* peel administered to 3-week-old chicks by oral gavage did not induce signs of toxicity [2].

When the tachyzoites multiplied in HFF host cells, the concentration required for 50% inhibition of *T. gondii* growth was lower for aqueous extract of *P. granatum* peel than for aqueous extract of *A. visnaga* seeds, while the percentage of annexin V+ tachyzoites was lower when treated with 5 µg/mL and 500 µg/mL of *P. granatum* extract than with the aqueous extract of *A. visnaga* seeds, suggesting distinct mechanisms of action in the two culture conditions. Indeed, *P. granatum* extract could also have an effect on host cells by inhibiting NF-κB activation, as it has been demonstrated in chondrocytes and microglial cells [3,22]. The plant extract could therefore inhibit the transcription of NF-κB-dependent anti-apoptotic genes and the subsequent extension of host cell lifespan, as described in fibroblasts infected with *T. gondii* tachyzoites [35].

In contrast to *A. visnaga*, numerous *in vitro* and *in vivo* studies have demonstrated the antioxidant properties of pomegranate [18]. An ethanolic extract of *P. granatum* peel was shown to be more effective than aqueous and acetonic extracts in scavenging the DPPH radical (50% activity at 7.36 ± 1.53 µg/mL, 11.3 ± 1.54 µg/mL and 13 ± 1.9 µg/mL, respectively) [5]. However, it is not clear how these concentrations were calculated. Indeed, in DPPH radical scavenging activity figure, the concentrations required to achieve 50% of the activity were between 30 and 60 µg/mL for the aqueous extract and above 120 µg/mL for the other two extracts [5]. In the literature, the antioxidant and antimicrobial activities of *P. granatum* have been attributed to tannins and phenols [18]. Since sterols were abundant in our aqueous extract of *A. visnaga* seeds, which has low anti-*Toxoplasma* activity, it is more likely that the gallic tannins of *P. granatum* were involved in the overall biological effects observed here.

The aqueous extract of cloves was antioxidant, efficient against *T. gondii* and not cytotoxic. Although phenols have been considered the main bioactive compounds of cloves [14], it is difficult to determine which classes of compounds were important for the anti-*Toxoplasma* and antioxidant activities of this aqueous extract. The active compounds of *S. aromaticum* flower buds extracted with the solvents dichloromethane, ethanol or hexane exhibited higher antioxidant activity than our aqueous extract with an oxidation inhibition of 42% to 93% at concentrations from 50 to 400 µg/mL [37]. The World Health Organization has recommended not to ingest more than 2.5 mg/kg/day of this spice, although only the toxicity of the essential oil and its main component eugenol and not of an aqueous extract has been studied [9]. Additional toxicity studies should be done to conclude.

In the case of plants, molecules belonging to different classes of compounds were certainly involved in the overall effect of the extracts. However, it is difficult to identify which ones were essential. In a previous study, we showed that an ethanolic extract and a hydro-ethanolic extract of teak bark (*Tectona grandis*) with equivalent contents of phenols, tannins and flavonoids had very different selectivity indices on *T. gondii* (10 and 0.3, respectively) [15]. On the contrary, an ethanolic and a hydro-ethanolic extract of teak leaves both gave a selectivity index lower than 1 (0.3 and 0.8, respectively) while the former contained more phenols and flavonoids and less tannins than the latter. In any case, the aim of our previous and current studies was not to identify a single active compound in order to purify or chemically synthesise it, but to underline the interest of using plant extracts containing several active compounds acting in association or synergy.

Even if an aqueous extract can be made in an artisanal way, an important parameter to consider for larger scale production is the extraction yield of the plant material. This ranged from 30% to 56% for the aqueous extracts (Table 2). The most promising candidate for the treatment of toxoplasmosis was the ethanolic extract of *A. visnaga*, but besides the fact that it requires a chemical to be made, the extraction yield was only 12%. It is important to note that *A. visnaga* is already marketed in many forms (capsule, tablet, granule, liquid, effervescent…) in European countries (France, Germany and Spain) and in Egypt [25]. These drugs are contraindicated during pregnancy [25], but it would be interesting to study their protective effect in clinical trials on ocular or cerebral toxoplasmosis. The second promising candidate of the present study was the aqueous extract of *P. granatum* peel. In a recent review [31], the authors highlighted the economic interest of exploiting this waste from the food industry. They also pointed out the possible differences in the contents of bioactive compounds depending on the methods used during the process, such as drying of the peel (air, microwave, oven, sun or vacuum) or the extraction technique itself (solvents, supercritical fluid, ultrasound, microwave or pressurized liquid). Pomegranate peel extract deserves further study in *in vivo* models of toxoplasmosis, with a view to its application in humans.

## Acknowledgments

We thank Prof. Khayati Najat (Faculty of Science Ain Chock, Hassan II University, Casablanca) for identifying plants. This work was supported by the University of Tours and the University of Casablanca.

